# *Trpm8* modulates thermogenesis in a sex-dependent manner without influencing cold-induced bone loss

**DOI:** 10.1101/2020.03.18.996694

**Authors:** Adriana Lelis Carvalho, Annika Treyball, Daniel J. Brooks, Samantha Costa, Ryan J. Neilson, Michaela R. Reagan, Mary L. Bouxsein, Katherine J. Motyl

**Author notes:** Grant Funding:* K01AR067858 to Katherine Motyl, PhD, Principal Investigator, P30GM106391 to Robert Friesel, PhD, Program Director, P30GM103392 to Don St. Germain, MD, Program Director, P20GM121301 to Lucy Liaw, PhD, Program Director, U54GM115516 to Clifford Rosen, MD, Program Director. Address correspondence to:* Katherine J. Motyl, 81 Research Drive, Scarborough, ME 04074.

## Abstract

*Trpm8* (transient receptor potential cation channel, subfamily M, member 8) is expressed by sensory neurons, and it is involved in the detection of cold temperatures. Increased TRPM8 activity triggers an increase in uncoupling protein 1 (*Ucp1*)-dependent brown adipose tissue (BAT) thermogenesis. Bone density and marrow adipose tissue are regulated by rodent housing temperature and brown adipose tissue, suggesting one mediator of this effect may be TRPM8. In order to test whether TRPM8 was involved in the co-regulation of thermogenesis and bone homeostasis, we examined the bone phenotypes of one-year-old *Trpm8* knockout mice (*Trpm8-KO*) and then exposed them to a 4-week cold temperature challenge. Male *Trpm8-KO* mice had lower bone mineral density than WT, with smaller bone size (femur length and cross-sectional area) being the most striking finding, and exhibited a delayed cold acclimation with increased BAT expression of *Ucp1, Dio2* and *Cidea* at the end of the study. In contrast to males, female *Trpm8-KO* mice had low vertebral bone microarchitectural parameters, but no genotype-specific alterations in body temperature. Interestingly, *Trpm8* was not required for cold-induced trabecular bone loss in either sex, but may be required for cold-induced changes in marrow adiposity in males. Furthermore, bone marrow adipose tissue accumulation in females was significantly blunted by *Trpm8* deletion. In summary, we identified sex differences in the role of TRPM8 in maintaining body temperature, bone density and marrow adipose tissue. Identifying mechanisms through which cold temperature and brown adipose tissue influence bone could help to ameliorate potential bone side effects of obesity treatments designed to stimulate thermogenesis.

## Introduction

Maintaining body temperature homeostasis involves both the regulation of heat loss and generation of heat (i.e. thermogenesis). Mammals require energy to generate and/or conserve heat when in temperatures below their thermoneutral zone, which is room temperature in humans, and warmer (28-32°C) in mice [1]. During adaptation to cooler temperatures, sympathetic nervous system (SNS) output increases, elevating energy expenditure, fat oxidation, and heat production by brown adipose tissue (BAT). Uncoupling protein 1 (*Ucp1*) in the inner mitochondrial membrane of BAT generates heat by uncoupling oxidative phosphorylation from ATP production [2,3]. In addition to this, cold exposure can also trigger white adipose tissue (WAT) beiging to enhance heat production [4]. Previous studies have demonstrated a relationship between SNS activity and the skeleton, revealing that sympathetic outflow is not only involved in the activation of BAT thermogenesis for thermoregulation purposes, but that sympathetic tone can also suppress bone formation by signaling through the β2-adrenergic receptor (β2AR) expressed by osteoblasts [11][12]. Marrow adipocytes and osteoblasts have a common mesenchymal precursor and are known to be elevated in many conditions of low bone density. However, the SNS suppresses both bone formation and bone marrow adipose tissue (BMAT), and a four-week cold exposure in on-year-old C3H/HeJ mice leads to reduced BMAT at the most active sites of bone remodeling [13]. In that study, however, bone parameters were largely unchanged, and there was no indication of whether C57BL/6J mice (a commonly used model for skeletal research) would have bone loss from cold exposure. A more recent study, however, showed that thermoneutral (32°C) housing in C57BL/6J mice exerted a protective effect on the rodent skeleton, and this was associated with reduced expression of the *Ucp1* in BAT [1]. These findings suggest that long-term cold-stimulus may be detrimental to bone density, but it is unclear what particular thermoregulatory pathways may promote bone and marrow adipose tissue loss.

Transient receptor potential melastatin 8 (TRPM8) is a Ca^2+^ permeable, non-selective cation channel expressed by temperature-sensing sensory neurons and is known to be the principal detector of innocuous cold [14]. In mice, TRPM8 is activated by a wide temperature range, approximately between 28°C to 8°C, and also by cooling agents (e.g. menthol and icilin) [15]. Previously, TRPM8 was primarily described to work as a peripheral thermal sensor; however, there is evidence that this cold-sensing channel is also involved in thermoregulation by promoting heat conservation and energy homeostasis [16]. Furthermore, TRPM8 is a potential target for obesity treatment due to its function in regulating energy metabolism [16–21]. TRPM8 has been found to trigger non-shivering thermogenesis by BAT and the beiging of WAT in mice [21][22] and humans [19]. Moreover, *Trpm8* knockout (*Trpm8*-KO) mice are hyperphagic and have reduced fat oxidation when housed at 21°C, promoting the development of obesity [16]. However, it is unknown whether TRPM8 has a role in regulating bone density *in vivo* and by inference, whether modulation of TRPM8 by obesity therapeutics might influence bone.

Thus, we conducted this study to determine whether the cold-sensing TRPM8 channel would be required for any cold-induced changes in the skeleton and bone marrow adipose tissue. We hypothesized that *Trpm8* deletion would impair body temperature regulation but may protect mice from cold-induced bone loss and marrow adipose tissue changes. We exposed one-year-old male and female WT and *Trpm8* knockout (KO) mice to a cold housing (4°C) treatment and examined body temperature, peripheral and marrow adipose tissue, bone microarchitecture and remodeling. We found that male *Tprm8-KO* mice had impaired temperature regulation while females did not. Male *Trpm8-KO* mice had a more overt low bone density phenotype, but females also had reduced microarchitectural parameters in cortical bone of the femur and trabecular bone of the vertebrae. Overall, cold temperature reduced trabecular bone volume fraction in males, while reducing cortical area fraction in females, but these effects were not dependent upon genotype. Surprisingly, bone marrow adiposity was lower in *Trpm8-KO* females compared to WT, but there was no striking effect of cold on either genotype or sex of mice, with the exception of an interaction suggesting TRPM8 may be require for BMAT loss after cold in males. There were, however, sex and genotype specific effects on the utilization of peripheral adipose tissue stores, suggesting TRPM8 regulates the energy demands of thermogenesis in a sex-dependent manner.

## Materials and Methods

*Wild-type* (WT) and *Trpm8-knockout* (*Trpm8-KO*) mice were purchased from The Jackson Laboratory (stock #008198). Mice were bred and housed in a barrier animal facility at Maine Medical Center Research Institute (MMCRI) on a 14-hr light and 10-hr dark cycle. Mice were aged to 52 weeks while being housed up to four animals per cage at 22°C. Mice were given water and regular chow *ad libitum*. All animal protocols in this study were approved by the Institutional Animal Care and Use Committee (IACUC) of MMCRI, an Association for Assessment and Accreditation of Laboratory Animal Care (AAALAC)-accredited facility. Euthanasia was performed using isoflurane followed by cervical dislocation or decapitation.

### Dual-energy X-ray absorptiometry (DXA)

All mice had body weight measured prior to DXA. Areal bone mineral content (aBMC), bone mineral density (aBMD), lean mass and fat mass measurements were performed on each mouse by a PIXImus dual energy X-ray densitometer (GE Lunar, GE Healthcare). The PIXImus was calibrated daily with a mouse phantom provided by the manufacturer. Mice were ventral side down with each limb and tail positioned away from the body. Full-body scans were obtained, and the head was excluded from analyses because of concentrated mineral content in skull and teeth. DXA data were processed and analyzed with Lunar PIXImus 2 (version 2.1) software [23]. Adiposity index was defined as fat mass divided by fat-free mass.

### Cold exposure Study

Fifty-two week-old WT and *Trpm8-KO* mice were separated into individual cages. Half of the mice were maintained at room temperature, while the other half were moved to a cold room with automatic lights set to a 14 hr light, 10 hr dark cycle, equivalent to the light schedule of room temperature mice. Light intensity was not measured, but was comparable to that in the barrier animal facility and the source of the light was at the ceiling. Cold-treated mice were acclimatized to 18°C for 1 week, then 4°C for 3 weeks. This method of cold-temperature exposure has been previously published to produce changes in marrow adipose tissue C3H/HeJ mice and in C57BL/6J mice as well as used to determine dependence of cold-acclimation on energy expenditure in mice [13][24]. Rectal temperature was measured daily in cold room mice and every 2 hours on the day of change to 4°C with a Type T thermocouple rectal probe (RET-3, Physitemp Instruments, Inc., Clifton, NJ, USA) with a MicroTherma 2T hand-held thermometer (ThermoWorks, Inc., Lindon, UT; cat. no. THS-227-193). Cages contained standard bedding. Mice were fed regular chow and water *ad libitum.* Environmental temperature and humidity was measured and recorded daily. Mice were monitored for signs of pain and distress twice per day and every 2 hours on the day of change to 4°C. We monitored and recorded any of the following signs: shivering, hunched posture, reduced mobility, ruffled fur and vocalization. Signs that were found included occasional shivering and ruffled fur, and these generally resolved immediately if not associated with low core temperature. If core temperature fell below 30°C, mice were immediately removed to room temperature and euthanized. No mice exhibited temperatures below this threshold or exhibited signs of distress during the transition from 18°C to 4°C. During the 4°C period, two mice (one male +/+ and one male −/−) exhibited a low body temperature and were immediately euthanized. No mice died spontaneously.

### Histology

Femur and all adipose tissue samples were placed in 10% neutral buffered formalin (Sigma-Aldrich) for 48 hr and then stored in 70% ethanol. Bones were decalcified with EDTA and bone and adipose tissue samples were paraffin embedded and tissues sections of 5 μm were deparaffinized and dehydrated in a graded series (100-70%) of ethanol washes and stained with hematoxylin and eosin (H&E). Photomicrographs of H&E stained tissues were taken at 4x magnification.

### Bone Marrow Adipocyte Quantification and Analysis

A blinded analysis of bone marrow (BM) adipocytes was performed on each histology slide using ImageJ software (https://imagej.nih.gov/ij/) [25,26]. A global scale was set at 1000 pixels/mm based on the microscope imaging objective and settings. The tissue area (T.Ar) was selected in the distal femoral metaphysis (0.1 mm below the epiphyseal growth plate) using the polygon tool to contain the BM, excluding cortical bone. One histological slide per distal femur was measured. The T.Ar was measured using ImageJ to find the total tissue area, which was used to normalize adipocyte numbers per T.Ar for each image. Color images were converted to 32-bit and then thresholded using the *Huang* filter at the automatic level determined by ImageJ [25,26]. [Note: thresholding an image converts it to black (255) and white (0) values [25,26]. Images were then processed with the *Despeckle* and *Adjustable Watershed* plugins. [*Despeckle* was used to automatically remove salt-and-pepper background noise that could cause inaccuracies with particle analysis, while the *Adjustable Watershed* ImageJ segmentation algorithm was used to systematically correct any adjacent adipocytes that had been merged into 1, back into 2 distinct objects. ImageJ typically applies a watershed segmentation threshold at a level of 0.5, but with the *Adjustable Watershed* the amount of segmentation can be changed to correct for closely packed adipocyte, tissue tears, or staining inconsistencies [25,26]. A blinded investigator set the *Adjustable Watershed* within the range of 0.3-0.8 to select the most appropriate segmentation threshold to apply to each image based solely on the quality of the individual image. Next, particle analysis for BM adipocytes was performed within the T.Ar using the *Analyze Particles* plugin. This plugin scans a selected area and measures objects that meet parameters based on area and circularity [25,26]. The particle circumference range was set to 50-2500 pixels (area results displayed in mm^2^ due to the set global scale) and the circularity was set at 0.60-1.00 (1.00 being a perfect circle). These parameters were determined through an iterative, trial and error analysis of different images until a universal measurement system was decided upon and then used for every image. After particle analysis, the *Result* and *Summary* pop-up windows provided the individual and average area (mm^2^) for each counted adipocyte within the T.Ar; these data were used to quantify adiposity in each sample (based on adipocyte number and average adipocyte area).

### Micro-computed tomography (μCT)

A high-resolution desktop micro-tomographic imaging system (μCT40, Scanco Medical AG, Brüttisellen, Switzerland) was used to assess trabecular bone architecture in the distal femoral metaphysis and L5 vertebral body and cortical bone morphology of the femoral mid-diaphysis. Scans were acquired using a 10 μm^3^ isotropic voxel size, 70 kVP, 114 μA, 200 ms integration time, and were subjected to Gaussian filtration and segmentation. Image acquisition and analysis protocols adhered to guidelines for the assessment of rodent bones by μCT [27]. In the femur, trabecular bone microarchitecture was evaluated in a 1500 μm (150 transverse slices) region beginning 200 μm superior to the peak of the growth plate and extending proximally. In the L5 vertebral body, trabecular bone was evaluated in a region beginning 100 μm inferior to the cranial end-plate and extending to 100 μm superior to the caudal end-plate. The trabecular bone regions were identified by manually contouring the endocortical region of the bone. Thresholds of 335 mgHA/cm^3^ and 385 mgHA/cm^3^ were used to segment bone from soft tissue in the femur and L5 vertebrae, respectively. The following architectural parameters were measured using the Scanco Trabecular Bone Morphometry evaluation script: trabecular bone volume fraction (Tb.BV/TV, %), t rabecular bone mineral density (Tb. BMD, mgHA/cm^3^), specific bone surface (BS/BV, mm^2^/mm^3^), trabecular thickness (Tb.Th, mm), trabecular number (Tb.N, mm^−1^), and trabecular separation (Tb.Sp, mm), connectivity density (Conn.D, 1/mm^3^). Cortical bone was assessed in 50 transverse μCT slices (500 μm long region) at the femoral mid-diaphysis and the region included the entire outer most edge of the cortex. Cortical bone was segmented using a fixed threshold of 708 mgHA/cm^3^. The following variables were computed: total cross-sectional area (bone + medullary area) (Tt.Ar, mm^2^), cortical bone area (Ct.Ar, mm^2^), medullary area (Ma.Ar, mm^2^), bone area fraction (Ct.Ar/Tt.Ar, %), cortical tissue mineral density (Ct.TMD, mgHA/cm^3^), cortical thickness (Ct.Th, mm), cortical porosity (%), as well as bone strength indicators such as maximum, minimum and polar moments of inertia (*I*_*max*_, *I*_*min*_, and pMOI, mm^4^), which describe the shape/distribution of cortical bone (larger values indicate a higher bending strength). Cortical bone images were taken of the mouse using the median total area value within each group. The scale for all of the images is 1 mm equals 315 pixels.

### Real-time PCR

Adipose tissues (brown and gonadal adipose tissue) were collected from WT and *Trpm8-KO* mice for RNA extraction under liquid nitrogen conditions. Total RNA was prepared using the standard TRIzol (Sigma-Aldrich) method for tissues. cDNA was generated using the High Capacity cDNA Reverse Transcriptase Kit (Applied Biosystems) according to the manufacturer’s instructions. mRNA expression analyses was carried out using an iQ SYBR Green Supermix with a BioRad^®^ Laboratories CFX Connect Real-time System thermal cycler and detection system. TATA binding protein 1 (Tbp1) was used as an internal standard control gene for all quantification [28]. Primers used were from Integrated DNA Tecnologies (IDT) (Coralvile, IA) [29–31] and Primer Design (Southampton, UK). All primer sequences are listed in Supplementary Table I.

### Statistical analysis

GraphPad Prism 8® software was used to visualize data and perform statistical analyses. Data are presented as mean ± standard error. Student’s t test or two-way ANOVA was performed with Holm-Sidak *post hoc* multiple comparison test as needed. *p*<0.05 was considered statistically significant.

## Results

Body composition and two-dimensional areal bone density (n=14-19/group) were assessed at 52 weeks of age in mice housed at standard room temperature (22°C) (Figure 1). Both male and female *Trpm8-KO* mice had reduced body mass compared to WT mice, which was in part due to a significant reduction in lean mass in males (Figure 1A,B). In both sexes, two-way ANOVA indicated that *Trpm8* deletion significantly affected fat mass and % body fat, but in pairwise comparisons, only female % body fat was lower after *Trpm8* deletion (Figure 1C,D). Significant sex and genotype effects were found in bone mineral density (BMD) and bone mineral content (BMC) parameters (Figure 1E,F). In the latter, there was a significant interaction between sex and genotype such that *Trpm8* deletion caused significantly lower bone mineral content in males, but not in females (Figure 1F).

**Figure 1.**
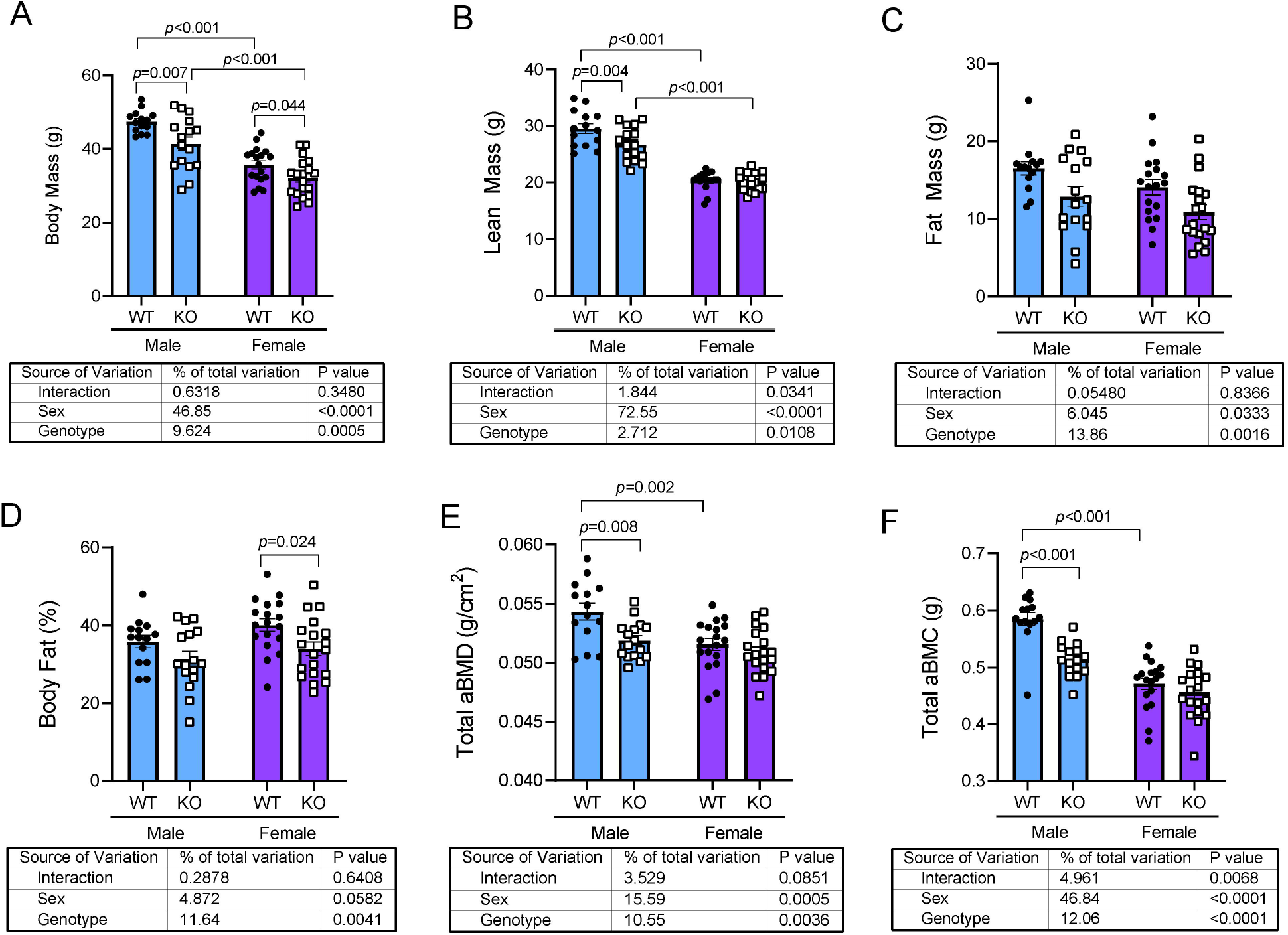
*Trpm8* deletion causes reduced body mass, lean mass and bone density in male mice. (A-D) Body composition in 52-week old male (blue) and female (purple), wildype (closed circles) and *Trpm8-KO* (open squares) mice was analyzed by DXA. (E,F) Areal total body bone mineral density (aBMD) and areal bone mineral content (BMC) were measured excluding the head. Data are expressed as individual points with bars representing mean ± SEM. Tables below each panel indicate results of the two-way ANOVA, with *p-*values for sex, genotype and their interaction. *p*-values above the bars represent results from Holm-Sidak *post hoc* test for pairwise significance.

Because significant changes in two-dimensional DXA data (aBMD and aBMC) are most likely to indicate alterations in cortical microarchitecture, we next examined whether *Trpm8* deletion led to changes in cortical bone by micro Computed Tomography (μCT) (Figure 2). Male *Trpm8-KO* mice had significantly lower femur total cross-sectional area (total area) and lower marrow area compared to male WT mice (Figure 2A-E). These differences were not significant in females. Polar moment of inertia (an indicator of strength), was dependent upon genotype (p=0.0479) but within sex pairwise comparisons did not reach statistical significance (Figure 2G). Other parameters, such as cortical tissue mineral density, cortical thickness and porosity were not dependent upon genotype (Figure 2F,H-J). However, femur length was shorter in male *Trpm8-KO* mice compared to WT (Figure 2K). Finally, to test whether changes in bone density were a result of altered trabecular microarchitecture, we performed μCT on the distal femur trabecular bone (Figure 3). Deletion of *Trpm8* did not influence volumetric trabecular architecture in male or female mice (Figure 3). Taken together, these results indicate smaller bone size in the male *Trpm8-KO* mice may have led to the reduced total body bone mineral content and density.

**Figure 2.**
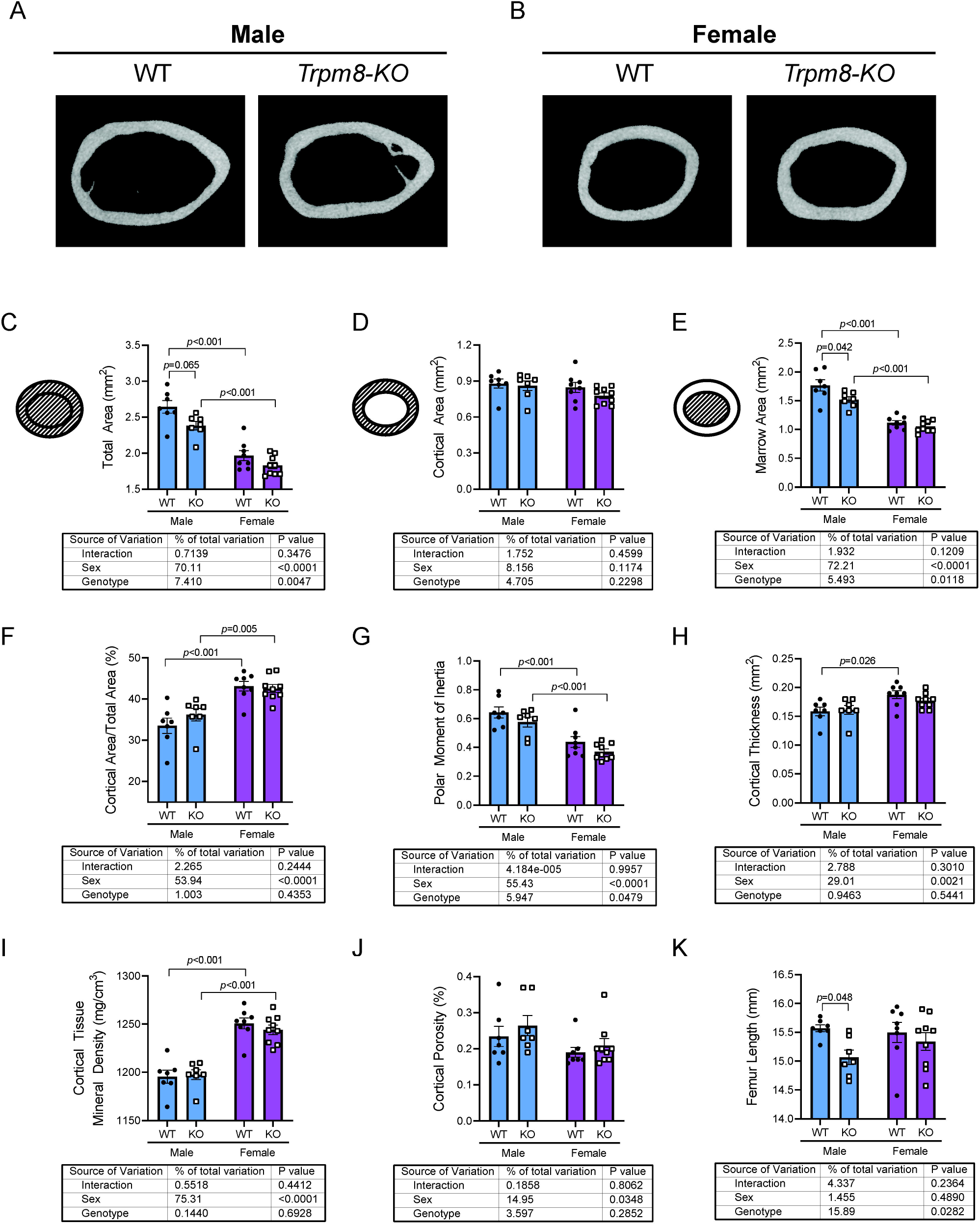
*Trpm8* deletion reduced femur size in male mice. Femurs from room temperature housed male and female *Trpm8-KO* and WT mice were scanned by μCT. (A,B) Representative images of the femur cross section from the midshaft, where cortical microarchitectural measurements were performed (C-J). Shaded areas of the pictograms in C-D represent the region of interest for each measurement. Data are expressed as individual points with bars representing mean ± SEM. Tables below each panel indicate results of the two-way ANOVA, with *p-*values for sex, genotype and their interaction. *p*-values above the bars represent results from Holm-Sidak *post hoc* test for pairwise significance.

**Figure 3.**
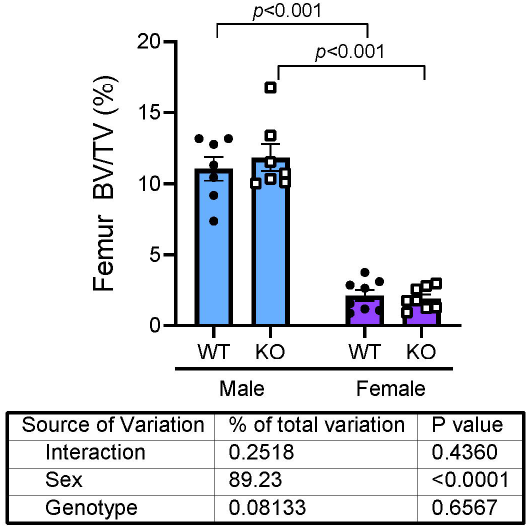
*Trpm8* deletion did not overtly impact trabecular microarchitecture of the distal femur. Bone volume fraction (BV/TV) of the trabecular bone of the distal femur was measured in male and female *Trpm8-KO* and WT mice. Data are expressed as individual points with bars representing mean ± SEM. Table below the panel indicate results of the two-way ANOVA, with *p-*values for sex, genotype and their interaction. *p*-values above the bars represent results from Holm-Sidak *post hoc* test for pairwise significance.

Previous studies have demonstrated that TRPM8 regulates brown adipose tissue function, and thermogenesis has been associated with changes in trabecular bone remodeling [12,22,23]. We next wanted to test, 1) whether a 4-week cold exposure would lead to trabecular bone loss and 2) whether TRPM8 was necessary for the adipose tissue and bone responses to cold (Figure 4A). Mice were separated into individual cages and then left at room temperature or moved to 18°C (Figure 4A). After one week, the temperature was changed from 18°C to 4°C. Male *Trpm8-KO* mice had a poorer response to the cold challenge than WT, with core temperature being lower in the KO (Figure 4B). Interestingly, the impact of housing temperature change on female mouse core temperature was not dependent upon genotype. By the end of week 2, male *Trpm8-KO* mice had acclimated to the cold exposure and reached core temperatures similar to those of male WT mice (Figure 4C) which was maintained until the endpoint of the study.

**Figure 4.**
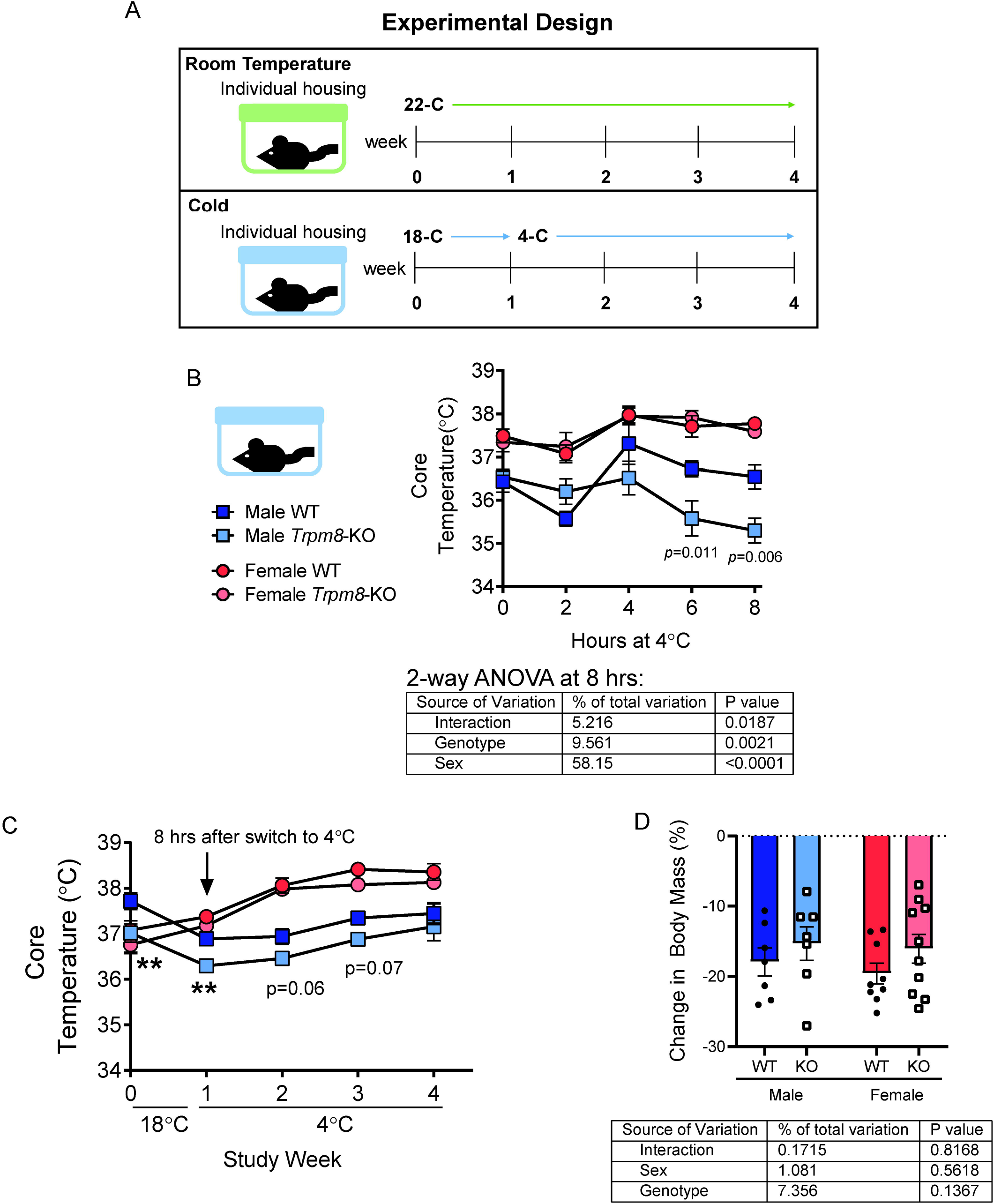
Male *Trpm8-KO* mice had lower core temperature during acclimatization to cold. (A) At 52 weeks of age, mice were separated into individual cages and half were kept at room temperature while the other half were moved to a cold room set at 18°C. After one week, the temperature was changed to 4°C and mice were maintained for 3 additional weeks. (B) During the temperature change from 18°C to 4°C, core temperature was measured every 2 hours. (C) Weekly core temperature, starting from time zero before mice were moved into the cold, and including the data from the 8 hour time point of the day the temperature changed from 18°C to 4°C. (D) Percent change in body weight. Temperature data points represent mean ± SEM. Body weight data are expressed as individual points with bars representing mean ± SEM. Table below panels B and D indicate results of the two-way ANOVA (performed using data from 8 hr time point in B), with *p-*values for sex, genotype and their interaction.

Previous studies had shown that impaired BAT function in response to cold was associated with bone loss. Therefore, we next wanted to identify changes in brown adipose tissue with cold treatment. Cold exposure induced BAT expansion in male WT and *Trpm8-KO* mice (p<0.05) (Figure 5A). In contrast, the expansion of BAT mass in female mice was observed only in the WT group (p=0.002) in response to cold, but not in female *Trpm8-KO* mice. In H&E stained sections of BAT, we observed larger lipid droplets in BAT from males of all genotypes and treatments, as well as in female *Trpm8-KO* mice at room temperature. Other female groups had less lipid accumulation, including in the female *Trpm8-KO* mice after cold. We also observed that cold treatment increased *Ucp1* gene expression in BAT from male WT mice, but the level of *Ucp1* expression was not different between cold treated WT and *Trpm8-KO* at either temperature. Overall, KO cold-treated mice had elevated expression of other fatty acid metabolism markers (*Acadl, Prdm16,* and *Pparc1α*) and thermogenic markers (*Cidea* and *Dio2*) (Figure 5C). Changes in BAT gene expression were less striking in females compared to males, with the notable exceptions of cold treatment inducing *Ucp1* to a greater degree in the KO females than in the WT females and suppressing *PParg2* in both genotypes (Figure 5C). Together, these results indicate *Trpm8* regulates the BAT response to temperature change in males and females.

**Figure 5.**
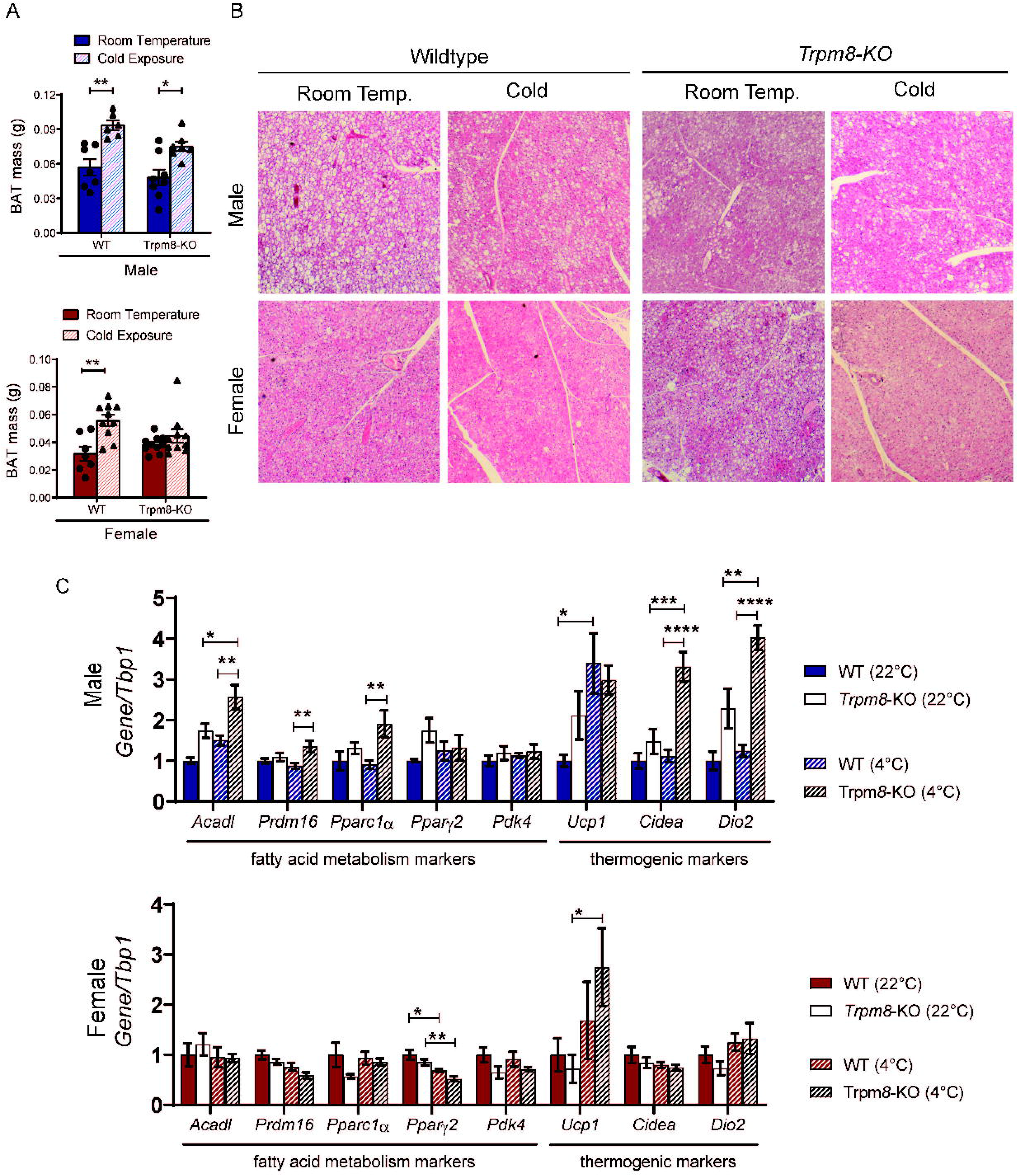
TRPM8 regulates the response of BAT to cold temperature. (A) BAT weights from male and female WT and *Trpm8-KO* housed at 22°C and 4°C. (B) Representative H&E stained image of BAT. N=5-8. (C) Gene expression of fatty acid metabolism and thermogenic markers in male (top) and female (bottom) BAT. Data presented as mean ± standard error. *p<0.05 and **p<0.01 by Holm-Sidak *post doc* test after a significant two-way ANOVA.

Although we did not find any significant differences in inguinal WAT weight (Figure S1), we did identify that gonadal adipose tissue (gWAT) mass was significantly lowered by cold treatment in male WT mice and in female *Trpm8-KO* mice (Figure 6A). H&E staining was suggestive of the presence of beige adipocytes after cold treatment in all sexes and genotypes except the *Trpm8-KO* cold treated males. The development of beige adipocytes was most striking in female *Trpm8-KO* mice, however, indicating a clear sex-difference in the way TRPM8 regulates the acclimatization to cold. These findings were supported by gene expression in gWAT in female mice, which showed, among other things, a 5 to 8-fold increase in *Ucp1* expression after cold, while the male mice only had a 2- and 4-fold increase in WT and KO, respectively, after cold treatment (Figure 6C). Similarly, female mice also had significant increases in *Acadl, Pparc1a, Pdk4, Cidea* and *Dio2* in the cold, whereas males did not have any other changes (Figure 6C). Taken together, these data indicate TRPM8 may be required for beiging of gWAT tissue in males but not females. Further, females may be more prone to generate beige adipocytes in gWAT than males, which could explain their advantage over males in adjusting to cold temperature.

**Figure 6.**
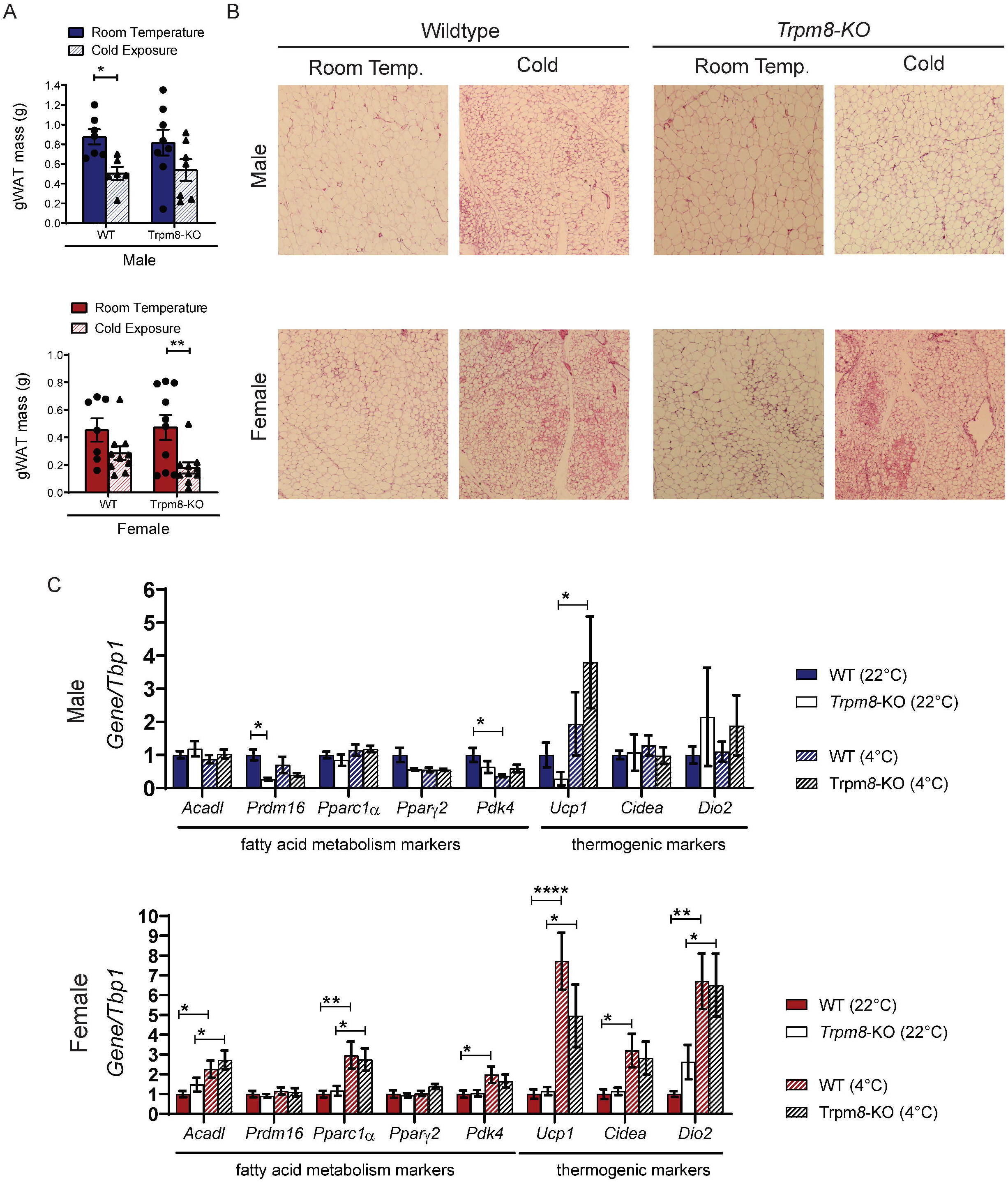
Loss of *Trpm8* prevented beige adipocyte formation in gonadal adipose tissue from male but not female mice. (A) gWAT weights from male and female WT and *Trpm8-KO* housed at 22°C and 4°C. (B) Representative H&E stained image of gWAT. (C) Gene expression of fatty acid metabolism and thermogenic markers in male (top) and female (bottom) gWAT. Data presented as mean ± standard error. *p<0.05 and **p<0.01 by Holm-Sidak *post doc* test after a significant two-way ANOVA.

Because altered thermogenesis has previously been associated with changes in bone homeostasis, we next examined bone microarchitectural phenotypes of WT and *Trpm8-KO* mice at room temperature and with cold exposure. In addition to cortical bone changes due to genotype and sex noted above in mice at room temperature (Figure 2), we determined that cold temperature caused trabecular bone loss (decreased BV/TV) in male mice of both gentoypes, in both the femur and the vertebrae (Table I). However, cold temperature treatment did not influence cortical bone in either genotype of male mice. In contrast to males, females had only had increased L5 trabecular BS/BV (indicative of trabecular thinning) in response to cold, but no changes in BV/TV in either the femur or the vertebral trabecular bone (Table II). Medullary area was increased with cold, leading to a reduction in cortical area fraction in females (Table II). However, there were no significant temperature by genotype interactions in trabecular or cortical bone parameters in either sex, suggesting bone loss from cold was not dependent upon TRPM8 in males or females (Tables I and II). Of note, female, but not male, mice also had reduced vertebral trabecular microarchitectural parameters after *Trpm8* deletion (Table II), but this was also not dependent upon housing temperature.

**Table I.** Femoral and Vertebral Bone outcomes assessed by μCT of Male wildtype (WT) and *Trpm8*-*Knockout* (*KO*) mice housed at room temperature or cold.

|  | Room temperature (22°C) |  | Cold (4°C) |  | *p*-value |  |  |
| --- | --- | --- | --- | --- | --- | --- | --- |
|  | WT | KO | WT | KO | genotype | temp | interaction |
| *Distal Femur Trabecular Bone* |  |  |  |  |  |  |  |
| BV/TV (%) | 11.1 ± 0.8 | 11.9 ± 0.9 | 8.9 ± 1.0 | 9.9 ± 1.1 | 0.36 | **0.04** | 0.91 |
| Conn.D (1/mm<sup>3</sup>) | 43 ± 7 | 58.5 ± 9.0 | 31.2 ± 5.8 | 43.1 ± 7.3 | 0.07 | 0.08 | 0.79 |
| Tb.N (1/mm) | 3.0 ± 0.1 | 3.2 ± 0.1 | 2.8 ± 0.1 | 3.0 ± 0.1 | 0.07 | 0.06 | 0.67 |
| Tb.Th (mm) | 0.059 ± 0.001 | 0.061 ± 0.003 | 0.058 ± 0.002 | 0.056 ± 0.002 | 0.95 | 0.15 | 0.19 |
| Tb.Sp (mm) | 0.323 ± 0.017 | 0.304 ± 0.007 | 0.347 ± 0.007 | 0.321 ± 0.011 | 0.07 | 0.09 | 0.78 |
| *Femur Midshaft Cortical Bone* |  |  |  |  |  |  |  |
| Ct.Th (mm) | 0.159 ± 0.007 | 0.161 ± 0.008 | 0.160 ± 0.006 | 0.154 ± 0.004 | 0.20 | 0.67 | 0.52 |
| Ct.Ar (mm<sup>2</sup>) | 0.88 ± 0.04 | 0.86 ± 0.04 | 0.88 ± 0.03 | 0.82 ± 0.03 | 0.26 | 0.49 | 0.52 |
| Ct.TMD (mgHA/cm<sup>3</sup>) | 1195 ± 7 | 1197 ± 5 | 1198 ± 8 | 1192 ± 5 | 0.73 | 0.79 | 0.50 |
| Ma.Ar (mm<sup>2</sup>) | 1.77 ± 0.10 | 1.52 ± 0.05 | 1.74 ± 0.11 | 1.54 ± 0.06 | **0.01** | 0.97 | 0.81 |
| Tt.Ar (mm<sup>2</sup>) | 2.64 ± 0.09 | 2.38 ± 0.06 | 2.62 ± 0.12 | 2.36 ± 0.07 | **0.005** | 0.77 | 0.99 |
| Ct.Ar/Tt.Ar (%) | 33.5 ± 1.9 | 36.2 ± 1.5 | 33.7 ± 1.4 | 34.8 ± 0.9 | 0.21 | 0.67 | 0.58 |
| Ct. Porosity (%) | 0.23 ± 0.03 | 0.26 ± 0.03 | 0.20 ± 0.01 | 0.23 ± 0.02 | 0.27 | 0.14 | 0.87 |
| pMOI (mm<sup>4</sup>) | 0.64 ± 0.04 | 0.58 ± 0.04 | 0.64 ± 0.04 | 0.54 ± 0.03 | **0.04** | 0.63 | 0.72 |
| Imax (mm<sup>4</sup>) | 0.42 ± 0.03 | 0.40 ± 0.03 | 0.42 ± 0.03 | 0.37 ± 0.03 | 0.19 | 0.53 | 0.67 |
| Imin (mm<sup>4</sup>) | 0.22 ± 0.01 | 0.18 ± 0.01* | 0.22 ± 0.02 | 0.18 ± 0.01* | **0.0009** | 0.89 | 0.89 |
| *L5 Vertebral Trabecular Bone* |  |  |  |  |  |  |  |
| BV/TV (%) | 21.6 ± 1.1 | 20.7 ± 1.8 | 18.7 ± 0.5 | 18.4 ± 0.7 | 0.59 | **0.04** | 0.78 |
| BS/BV (mm<sup>2</sup>/mm<sup>3</sup>) | 43.9 ± 1.5 | 44.5 ± 1.9 | 46.2 ± 0.4 | 46.9 ± 0.3 | 0.62 | 0.08 | 0.95 |
| BMD (mgHA/cm<sup>3</sup>) | 226.4 ± 10.0 | 217.3 ± 16.1 | 198.2 ± 4.7 | 197.6 ± 5.9 | 0.65 | **0.03** | 0.69 |
| Conn.D (1/mm<sup>3</sup>) | 127 ± 11 | 115 ± 8 | 113 ± 5 | 119 ± 8 | 0.73 | 0.53 | 0.32 |
| Tb.N (1/mm) | 4.4 ± 0.1 | 4.3 ± 0.1 | 4.1 ± 0.1 | 4.2 ± 0.1 | 0.94 | 0.11 | 0.43 |
| Tb.Th (mm) | 0.050 ± 0.001 | 0.053 ± 0.002 | 0.050 ± 0.003 | 0.053 ± 0.002 | 0.10 | 0.96 | 0.96 |
| Tb.Sp (mm) | 0.215 ± 0.007 | 0.223 ± 0.009 | 0.236 ± 0.005 | 0.230 ± 0.009 | 0.89 | 0.08 | 0.37 |

\*p<0.05 compared to WT mice group. N=6–7. *Abbreviations:* BV/TV, bone volume fraction; Conn.D, connectivity density; Tb.N, trabecular number; Tb.Th, trabecular thickness; Tb.Sp, trabecular separation; Ct.Th, average cortical thickness; Ct.Ar, cortical bone area; Ct.TMD, cortical tissue mineral density; M.Ar, marrow area; T.Ar, total cross-sectional area; Ct.Ar/T.Ar, cortical area fraction; Ct.Porosity, cortical porosity; pMOI, polar moment of inertia; I<sub>max</sub>, maximum moment of inertia; I<sub>min</sub>, minimum moment of inertia; BMD, bone mineral density; BS/BV, specific bone surface.

**Table II.**
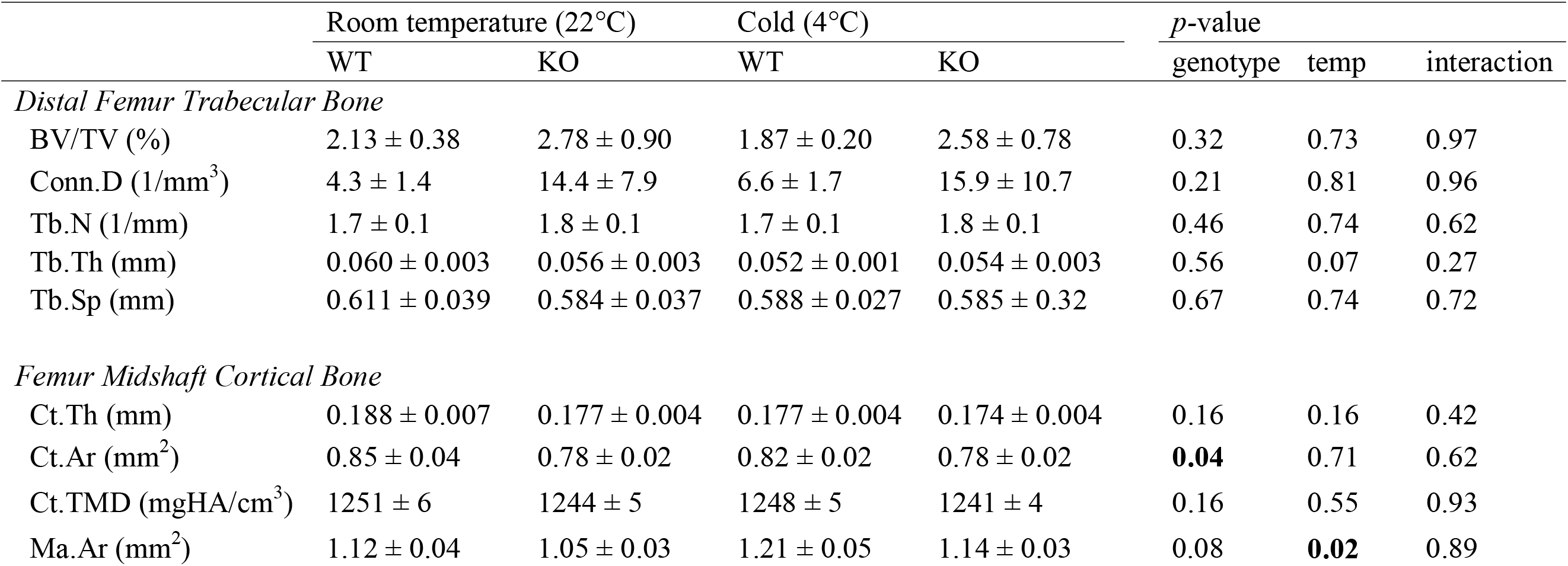
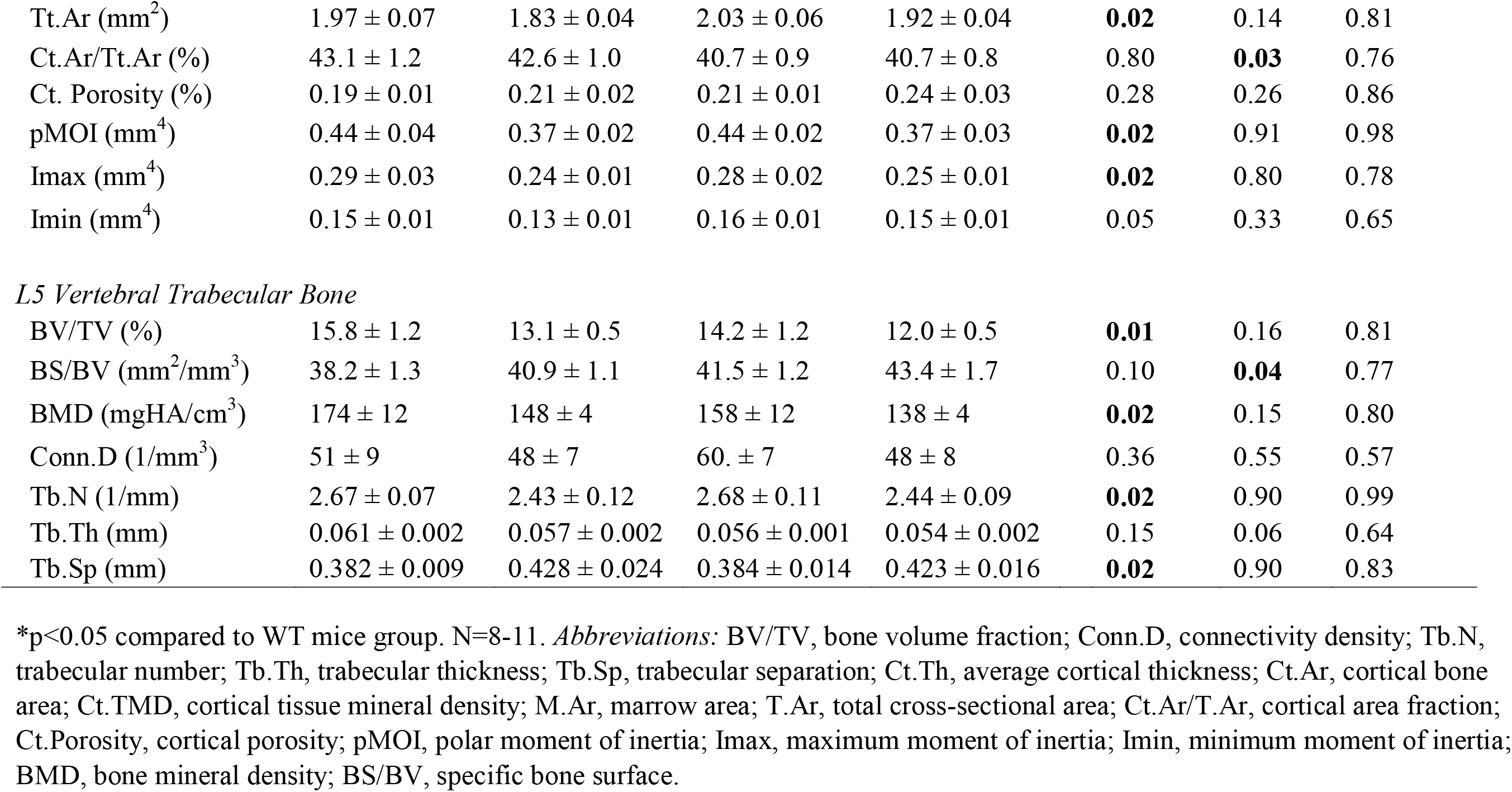
Femoral and Vertebral Bone outcomes assessed by μCT of Female wildtype (WT) and *Trpm8*-*Knockout* (*KO*) mice housed at room temperature or cold.

|  | Room temperature (22°C) |  | Cold (4°C) |  | *p*-value |  |  |
| --- | --- | --- | --- | --- | --- | --- | --- |
|  | WT | KO | WT | KO | genotype | temp | interaction |
| *Distal Femur Trabecular Bone* |  |  |  |  |  |  |  |
| BV/TV (%) | 2.13 ± 0.38 | 2.78 ± 0.90 | 1.87 ± 0.20 | 2.58 ± 0.78 | 0.32 | 0.73 | 0.97 |
| Conn.D (1/mm<sup>3</sup>) | 4.3 ± 1.4 | 14.4 ± 7.9 | 6.6 ± 1.7 | 15.9 ± 10.7 | 0.21 | 0.81 | 0.96 |
| Tb.N (1/mm) | 1.7 ± 0.1 | 1.8 ± 0.1 | 1.7 ± 0.1 | 1.8 ± 0.1 | 0.46 | 0.74 | 0.62 |
| Tb.Th (mm) | 0.060 ± 0.003 | 0.056 ± 0.003 | 0.052 ± 0.001 | 0.054 ± 0.003 | 0.56 | 0.07 | 0.27 |
| Tb.Sp (mm) | 0.611 ± 0.039 | 0.584 ± 0.037 | 0.588 ± 0.027 | 0.585 ± 0.32 | 0.67 | 0.74 | 0.72 |
| *Femur Midshaft Cortical Bone* |  |  |  |  |  |  |  |
| Ct.Th (mm) | 0.188 ± 0.007 | 0.177 ± 0.004 | 0.177 ± 0.004 | 0.174 ± 0.004 | 0.16 | 0.16 | 0.42 |
| Ct.Ar (mm<sup>2</sup>) | 0.85 ± 0.04 | 0.78 ± 0.02 | 0.82 ± 0.02 | 0.78 ± 0.02 | **0.04** | 0.71 | 0.62 |
| Ct.TMD (mgHA/cm<sup>3</sup>) | 1251 ± 6 | 1244 ± 5 | 1248 ± 5 | 1241 ± 4 | 0.16 | 0.55 | 0.93 |
| Ma.Ar (mm<sup>2</sup>) | 1.12 ± 0.04 | 1.05 ± 0.03 | 1.21 ± 0.05 | 1.14 ± 0.03 | 0.08 | **0.02** | 0.89 |
| Tt.Ar (mm<sup>2</sup>) | 1.97 ± 0.07 | 1.83 ± 0.04 | 2.03 ± 0.06 | 1.92 ± 0.04 | **0.02** | 0.14 | 0.81 |
| Ct.Ar/Tt.Ar (%) | 43.1 ± 1.2 | 42.6 ± 1.0 | 40.7 ± 0.9 | 40.7 ± 0.8 | 0.80 | **0.03** | 0.76 |
| Ct. Porosity (%) | 0.19 ± 0.01 | 0.21 ± 0.02 | 0.21 ± 0.01 | 0.24 ± 0.03 | 0.28 | 0.26 | 0.86 |
| pMOI (mm<sup>4</sup>) | 0.44 ± 0.04 | 0.37 ± 0.02 | 0.44 ± 0.02 | 0.37 ± 0.03 | **0.02** | 0.91 | 0.98 |
| Imax (mm<sup>4</sup>) | 0.29 ± 0.03 | 0.24 ± 0.01 | 0.28 ± 0.02 | 0.25 ± 0.01 | **0.02** | 0.80 | 0.78 |
| Imin (mm<sup>4</sup>) | 0.15 ± 0.01 | 0.13 ± 0.01 | 0.16 ± 0.01 | 0.15 ± 0.01 | 0.05 | 0.33 | 0.65 |
| *L5 Vertebral Trabecular Bone* |  |  |  |  |  |  |  |
| BV/TV (%) | 15.8 ± 1.2 | 13.1 ± 0.5 | 14.2 ± 1.2 | 12.0 ± 0.5 | **0.01** | 0.16 | 0.81 |
| BS/BV (mm<sup>2</sup>/mm<sup>3</sup>) | 38.2 ± 1.3 | 40.9 ± 1.1 | 41.5 ± 1.2 | 43.4 ± 1.7 | 0.10 | **0.04** | 0.77 |
| BMD (mgHA/cm<sup>3</sup>) | 174 ± 12 | 148 ± 4 | 158 ± 12 | 138 ± 4 | **0.02** | 0.15 | 0.80 |
| Conn.D (1/mm<sup>3</sup>) | 51 ± 9 | 48 ± 7 | 60. ± 7 | 48 ± 8 | 0.36 | 0.55 | 0.57 |
| Tb.N (1/mm) | 2.67 ± 0.07 | 2.43 ± 0.12 | 2.68 ± 0.11 | 2.44 ± 0.09 | **0.02** | 0.90 | 0.99 |
| Tb.Th (mm) | 0.061 ± 0.002 | 0.057 ± 0.002 | 0.056 ± 0.001 | 0.054 ± 0.002 | 0.15 | 0.06 | 0.64 |
| Tb.Sp (mm) | 0.382 ± 0.009 | 0.428 ± 0.024 | 0.384 ± 0.014 | 0.423 ± 0.016 | **0.02** | 0.90 | 0.83 |

\*p<0.05 compared to WT mice group. N=8-11. *Abbreviations:* BV/TV, bone volume fraction; Conn.D, connectivity density; Tb.N, trabecular number; Tb.Th, trabecular thickness; Tb.Sp, trabecular separation; Ct.Th, average cortical thickness; Ct.Ar, cortical bone area; Ct.TMD, cortical tissue mineral density; M.Ar, marrow area; T.Ar, total cross-sectional area; Ct.Ar/T.Ar, cortical area fraction; Ct.Porosity, cortical porosity; pMOI, polar moment of inertia; Imax, maximum moment of inertia; Imin, minimum moment of inertia; BMD, bone mineral density; BS/BV, specific bone surface.

Finally, we also evaluated whether cold exposure and/or loss of *Trpm8* would affect the adiposity of the bone marrow (Figure 7). In general, bone marrow adipose tissue (BMAT) is limited in male mice compared to females (Figure 7A-G). In male mice, we only found a significant interaction between genotype and housing temperature (p=0.045) in the % total adiposity, suggesting *Trpm8-KO* males do not lose marrow adipose tissue at cold temperatures, while WT mice do (Figure 7D). In female mice, there was a significant effect of *Trpm8* deletion on adipocyte number (Figure 7E) and % adiposity (Figure 7G), but not adipocyte size. In contrast to males, there was no clear impact of housing temperature on bone marrow adipose tissue females.

**Figure 7.**
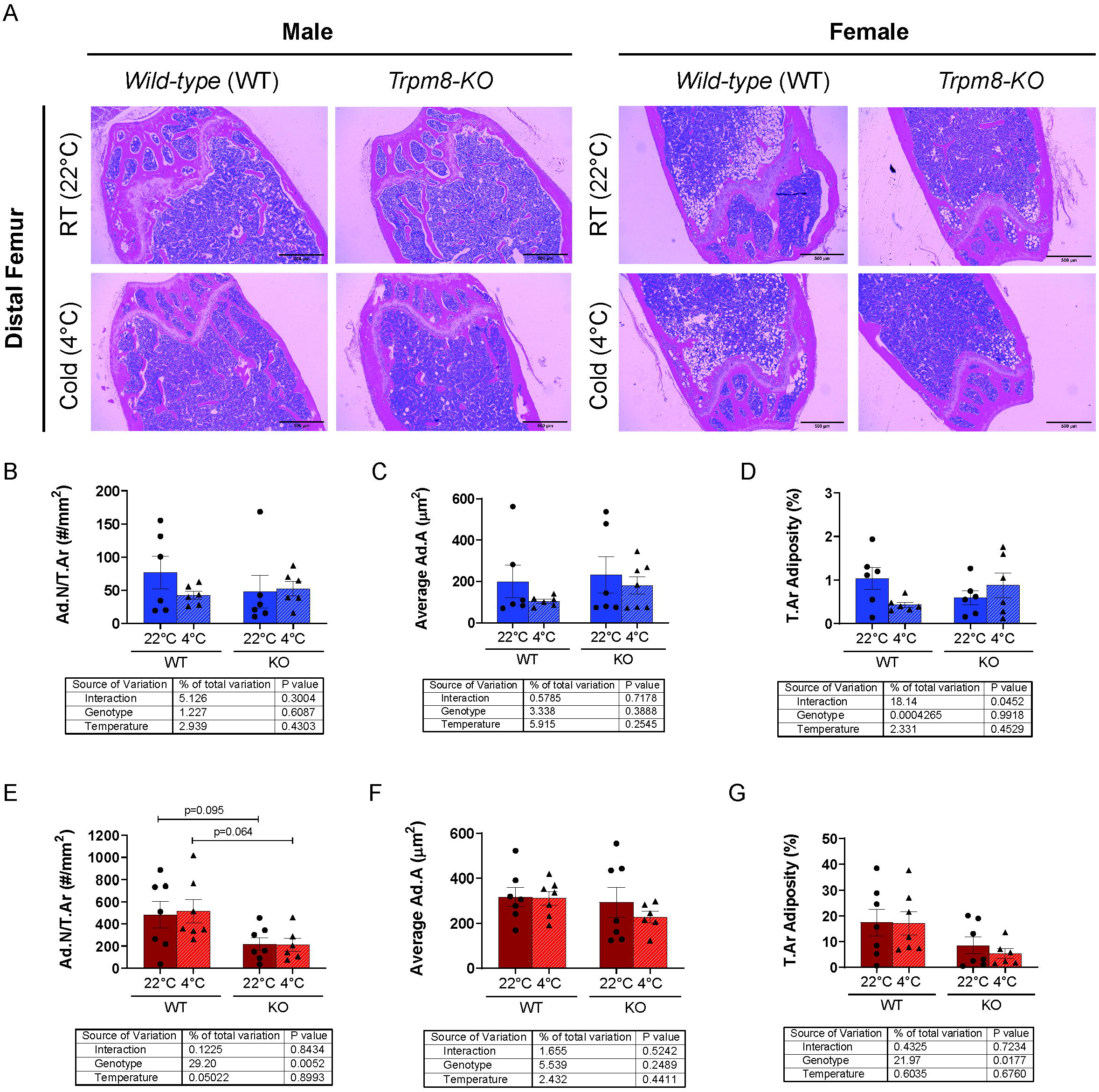
*Trpm8* deletion suppressed marrow fat in female mice. (A and B) Representative μCT images of cortical bone from male and female WT and *Trpm8-KO* mice housed at standard room temperature (RT) (22°C) and cold temperature (4°C). (C and D) Quantification of adipocyte numbers (Ad.N/T.Ar), average adipocyte size (Ad.A), and % adiposity (T.Ar Adiposity) within the bone marrow of male and female WT and *Trpm8-KO* mice housed at 22°C and 4°C. (E and F) Representative H&E stained image of distal femur area from male and female WT and *Trpm8-KO* mice housed at 22°C and 4°C (scale bar: 500 μm; magnification: 4x). Data presented as mean ± standard error.

## Discussion

In this study, we determined that the impact of *Trpm8* deletion on bone, body temperature and adipose tissue was dependent upon sex and housing temperature (Table IV summary). Our original hypothesis was that *Trpm8* deletion would impair body temperature regulation, but would also protect against cold-induced changes in bone and marrow adipose tissue. Males, but not females, had impaired temperature regulation after *Trpm8* deletion. However, cold housing led to trabecular bone loss in males and cortical bone loss in females, independent of genotype, which is consistent with previous studies showing that cold exposure can negatively impact the skeleton [1,12]. It has also previously been shown that cold exposure is associated with reduced BMAT [32], and our findings indicate that in males this may be dependent upon *Trpm8* (Figure 7). On the other hand, female BMAT was highly dependent upon *Trpm8* but not dependent upon housing temperature, pointing toward potentially diverse function of TRPM8 in regulating BMAT levels and response to stimuli. Despite the lack of any temperature by genotype interaction effects on bone parameters, it is worth noting that male KO mice, with impaired temperature regulation, also had smaller bones, while female KO mice had normal temperature regulation had normal bone size.

**Table IV.** Summary of results.

|  | **Males** |  | **Females** |  |
| --- | --- | --- | --- | --- |
|  | *Trpm8* Deletion | Cold Housing | *Trpm8* Deletion | Cold Housing |
| **Body Temperature** | Sex by genotype interaction – temperature reduced in male KOs but not female KOs in cold |  |  |  |
| **Bone Size** | decreased | uc | uc | uc |
| **Cortical Bone** | uc | uc | uc | decreased |
| **Trabecular Bone** | uc | decreased | decreased (L5 only) | uc |
| **Marrow Adiposity** | Genotype by temp interaction |  | decreased | uc |

*Abbreviation:* uc, unchanged.

A bone phenotype in *Trpm8-KO* mice has not been previously described. Bone size was smaller in male *Trpm8-*KO mice compared to WT, but this difference did not occur in females. Furthermore, female mice had significant trabecular bone loss in the vertebrae but not the femur, indicating potential site-specific effects of *Trpm8* deletion on bone. Recently, Iwaniec et al. (2016) highlighted that the current standard room temperature housing (22°C) can already be considered a cold environment to mice and that it was associated with bone loss in growing females compared to thermoneutral housing [1]. A *Trpm8* functional cold circuit is established after birth around postnatal day 14 [33]. Others have found that *Trpm8* is a regulator of energy metabolism and thermogenesis [16,19,21], and our findings are consistent: showing that its deletion can cause an increase in the expression of thermogenic markers, gWAT fat pad reduction and weight loss. This suggests that the absence of *Trpm8* during the developmental phase might be associated with defective cold circuits which partially fail to sustain optimal body temperature, as previously described [16]. Thus, it is plausible that the lack of *Trpm8* during growth causes a shift towards energy utilization for heat generation, decreasing ATP availability for bone cells, leading to altered bone density. However, the mechanism of low bone mass in *Trpm8-KO* mice, and the relative contributions of bone forming osteoblasts and bone resorbing osteoclasts, are yet to be determined. Some *in vitro* evidence suggests human osteoblast-like cells and murine osteoblast cells express *Trpm8,* but the functional impact on bone is unclear [34]. Although the deletion of other *Trp* channels, such as *Trpv1* [35] and *Trpv5* [36], has been reported to induce bone loss in animal models, there is a lack of data exploring the physiological role of TRPM8 in skeletal homeostasis.

One interesting finding was that *Trpm8* deletion in females caused decreased BMAT content. Aging generally leads to accumulation of bone marrow adipose tissue in the distal femur and proximal tibia in mice, but the mechanisms and consequences of this accumulation remain an area of active investigation. Both osteoblasts and adipocytes arise from a shared mesenchymal precursor, thus one hypothesis is that situations of bone loss may lead to marrow adiposity by shunting cells away from the osteoblast lineage to the adipocyte lineage (as in calorie restriction, PPARg2 inhibition, and aging). However, the alternate, shifting from marrow adipose lineage to osteoblast lineage, does not seem likely, since female *Trpm8-KO* mice had indices of reduced bone density in concert with low marrow adiposity. In other situations, like bone loss from the atypical antipsychotic drug risperidone, elevated SNS activity leads to a reduction in both osteoblast-mediated bone formation and marrow adiposity as well as increased expression of *Ucp1* in BAT [23]. Recently, Wee et al., (2019) elegantly demonstrated that iWAT and BMAT shared sympathetic innervation using retroviral nerve tracing [37]. They revealed that leptin-responsive regions such as paraventricular hypothalamus, arcuate nucleus, dorsomedial hypothalamus, ventromedial hypothalamus, and area postrema were labelled after viral tracing from iWAT, bone marrow, and BMAT [37]. However, it remains unclear if sex-differences exist in neural regulation of these tissues, and if gWAT would be regulated by similar common sympathetic pathways. Taken together, these findings suggest elevated SNS activity to stimulate thermogenesis could be responsible for bone loss and reduced marrow adiposity with *Trpm8* deletion.

Investigations into sex differences in the thermogenic response to *Trpm8* deletion have been limited. Clearly, *Trpm8* is important for the detection of environmental cooling [14,15,38] and it has also been implicated in BAT thermogenesis and thermoregulatory processes [16,21][22]. Early publications on *Trpm8* had reported no differences in core temperature associated with its global deletion in mice [38]. Ma et al. (2012) demonstrated that menthol activates Ucp-1 dependent thermogenesis and increases core temperature in WT mice, but that when TRPM8 is deleted, this activation is absent [22]. In contrast, our studies use cold-temperature to induce thermogenesis, and show a partial dependence on TRPM8. However, our findings also indicate that other pathways may compensate for the loss of TRPM8, especially in the females. Specifically, *Trpm8-KO* male mice struggled to increase body temperature to the level of WT in a cold environment (of which, menthol treatment is partly mimicking). However, the study by Ma, et al. (2012) did not directly compare WT to KO body temperature, so it is difficult to align perfectly with our findings. Recently, Reimúndez et al. (2018) demonstrated that *Trpm8* is required for thermoregulation in male mice when housed in colder environments, which corroborates our findings [16]. Interestingly, Reimúndez et al. (2018) also demonstrated that this difficulty in maintaining core temperature was associated with greater tail vasodilation and heat loss in male mice with global deletion of *Trpm8*, indicating that this cold-sensing channel has an important role in regulating both thermogenesis and heat dissipation [16]. However, female core temperature status was not investigated in any of these studies.

In contrast, one strength of our study is the comprehensive analyses of mice of both sexes. Cooling sensitivity is the first step to activate the cold response neuronal mechanism that modulates lipolysis and energy expenditure [39]. In our study, we observed that cold-induced weight loss occurred in both sexes, irrespective of the genotype. However, there were *Trpm8*- and sex-differences among the various adipose tissues’ response to housing temperature. Male mice housed at 4°C exhibited BAT expansion with fewer lipid droplets and increased *Ucp1* expression in WT, which is indicative of thermogenesis activity. However, this difference did appear to be blunted in KO mice (Figure 5). Moraes et al. (2017) found reduced *Ucp1* and *Pparg* expression at ZT2 and ZT14 in male TRPM8-KO mice ranging from 3-6 months of age, compared to ours that were 12 months [20]. Although we did not see lower *Ucp1* or *Pparg* in our room temperature mice, we expect that this difference might be due to the age of the mice or the time of the day that our tissues were collected ZT6-ZT10. However, our overall finding that thermogenesis is impaired in males is consistent with Moraes et al. finding of reduced *Ucp1* [20]. Further, we found that cold housing in male KO, but not WT, mice induced BAT expression of other fatty acid metabolism markers (*Acadl, Prdm16,* and *Pgc1α*) as well as a robust expression of other thermogenic markers such as *Cidea* and *Dio2* in BAT, which may have been compensatory for a lack of *Ucp1* induction compared to room temperature.

In females, we found that *Trpm8* deletion led to an increase in *Ucp1* expression and reduction in lipid droplets in BAT during cold exposure, indicating TRPM8 was not required for the response. Furthermore, cold reduced gWAT mass in female mice from both genotypes, in addition to increasing expression of fatty acid metabolism (*Acadl, Pgc1α, Pdk4*) and thermogenic (*Ucp1, Cidea, Dio2*) markers. It should be considered that estrogen status may play a role in sex-differences in thermogenesis and WAT beiging found in the present study. Several studies observed that the beiging process does take place in gWAT in females [40,41]. Increased expression of beiging markers such as *Ucp1*, *Pgc1α*, *Elovl3*, *Cidea* was observed in gWAT after either 10 days of cold exposure [41] or β3-adrenergic receptor agonist treatment [40] in female mice. Kim et al. (2016) demonstrated that gWAT in females is more susceptible to the beiging process than in males, due to more extensive sympathetic innervation inducing brown adipogenesis [40]. Also, the authors showed that sympathetic innervation of gWAT is sex-hormone dependent, as reduced tyrosine hydroxylase protein levels and lack of beiging of gWAT was found after chemical ovarian ablation [40]. Indeed, lipolysis can be directly modulated by estrogen through estrogen receptor alpha (ERα) signaling in BAT and WAT adipocytes [42]. Estrogen also has been described to play a critical role in cold sensitivity and hyperalgesia [43,44], which are also functions thought to be regulated by *Trpm8* [45,46]. However, interestingly, a previous preclinical study observed that *Trpm8* expression was upregulated in the skin of ovariectomized rats compared to sham control rats [47] and β-estradiol treatment might reduce *Trpm8* overexpression in the skin of ovariectomized rats [48]. These findings support our data that suggest regulation of energy metabolism in female mice is independent of *Trpm8* cold-sensing function and estrogen might be a more relevant factor to normal thermoregulation in 26 females. It is unclear however, whether ovariectomized mice, or mice aged to natural amenorrhea, would have a response to cold that is modified by *Trpm8* deletion.

The global deletion of *Trpm8* may have led to compensatory changes in other thermogenic mechanisms, which may have been sex-dependent. In addition to TRPM8, there is another cold-activated *Trp* channel known as transient receptor potential ankyrin 1 (TRPA1) which is considered to be a noxious cold receptor [49]. Interestingly, TRPM8 and TRPA1 appear to function in both complementary and synergistic fashion to detect environmental innocuous and noxious cold [49]. Winter et al. (2017) found that the lack of *Trpm8* and *Trpa1* led to a striking reduction in cold avoidance behavior in mice [49]. However, an acute cold avoidance response was still observed in *Trpm8-KO* mice with functional TRPA1 when exposed to a temperature range between 28°C and 15°C [49]. This suggests that TRPA1 can only partially compensate for the absence of TRPM8 [49]. When cold-defense mechanisms, such as thermogenesis and vascular tone, fail to sustain an optimal body temperature, a shivering response is triggered to increase heat generation [50]. In our study, only occasional shivering-like behaviors were observed in some mice. These resolved quickly and we did not note any differences in the genotypes of the mice that shivered. Previous studies found that *Trpm8* is not required to induce shivering and tachycardia during cold challenge [50]. Instead, shivering is found to be regulated by TRPV1 [50], which once more indicates TRPM8 is a major regulator of non-shivering thermogenesis.

In summary, our findings support that *Trpm8* is important for body temperature maintenance in male mice and its absence triggers a compensatory response to counteract reduction in core temperature. Moreover, *Trpm8* does not appear to regulate the bone response to colder environments, where it is presumed that the sympathetic nervous system-mediated stress response and/or diversion of energy away from bone contributes to bone loss. Interestingly, deletion of *Trpm8* was associated with bone size and shape differences in male mice as well. Therefore, although any bone loss associated with acute cold housing may be independent of TRPM8, low bone mass found in *Trpm8-*KO mice compared to WT, more striking in males, indicates that the functions of TRPM8 could contribute to bone modeling and remodeling throughout lifespan.

## Limitations

Some limitations of this study do exist. First, addition of a thermoneutral group would potentially exacerbate differences observed, and could provide additional insight into whether *Trpm8* regulates bone and marrow adiposity independent of cold temperature sensation and subsequent physiologic response. We regret that we did not have the capability to age mice to 52 weeks of age in a thermoneutral (~28-30°C) rodent incubator when this study was first designed. Next, because the room temperature mice were housed in a separate room from the cold-treated, we cannot exclude the possibility that the additional noise of the cold room contributed to the phenotype. There is no known association between noise stress and changes in brown adipose tissue, but there are some studies examining the impact of chronic mild stress (CMS), of which one of the stressors is noise, on bone density. One study found that the CMS caused bone loss, and this could be blocked by propranolol, which suggests the SNS is involved [54]. However, other stressors were involved in that study, including water deprivation, continuous overnight illumination, stroboscopic illumination, cage tilt, and time in a soiled cage. Thus, it is unclear whether noise alone would have influenced bone microarchitecture. Although the sound was greater in the cold room in our study, it was consistent over time, and the fact remains that compared to WT males, and females of both genotypes all treated in the cold, the body temperature in the male KO mice had a delayed response to cold. Other key findings, like *Trpm8-KO* mediated bone loss, were not dependent upon housing conditions. Light intensity was not measured, but was similar and from the ceiling in both rooms. Furthermore, additional N would have helped to reduce biological variability, which was high in some of the secondary outcomes of interest, such as inguinal WAT mass and expression of *Ucp1* in both BAT and gWAT. Although differences were not detected in iWAT mass, it is possible that histology and mRNA analyses of this tissue would have provided additional valuable information. However, iWAT samples were not saved to analyze at a later time point. Additionally, examining the thermogenic response at the time point where we observed major differences in body temperature in male mice exposed to cold would have provided better insight into mechanism. However, analysis of bone defects after cold were the primary outcomes of interest, and these would not have been apparent at such an early time point.

Despite these limitations, the current data provide new insight into our understanding of the role of *Trpm8* in the regulation of thermogenesis and skeletal homeostasis and demonstrate important and unexpected sex-differences in these processes. This work suggests that in males in particular, TRPM8 is necessary for normal thermoregulation and bone density, while in females, TRPM8 regulates marrow adiposity.

---

**Supplemental Table I.**
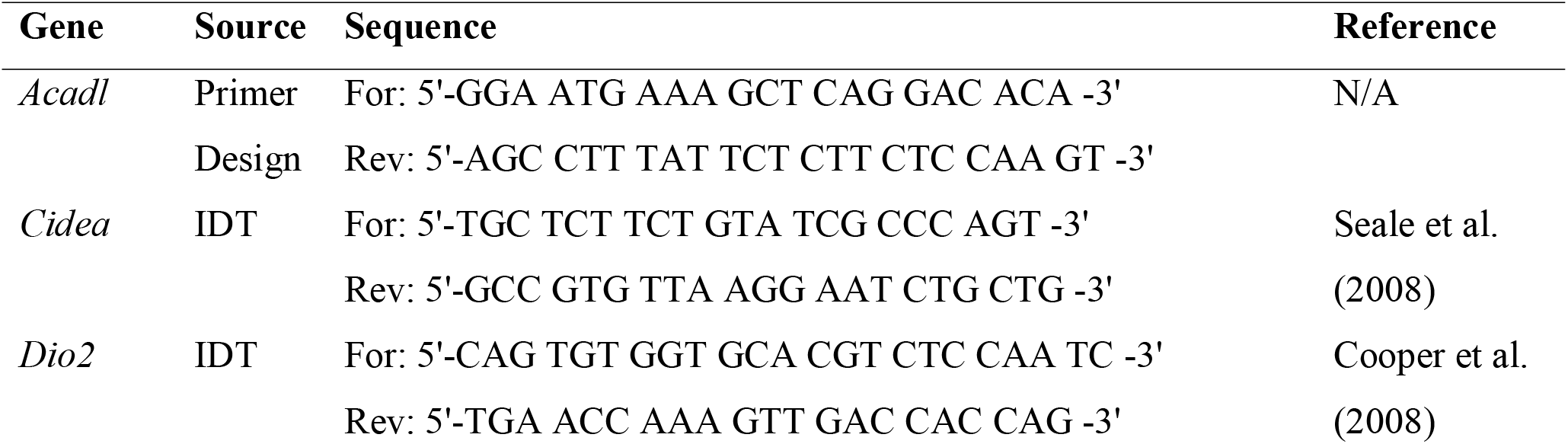
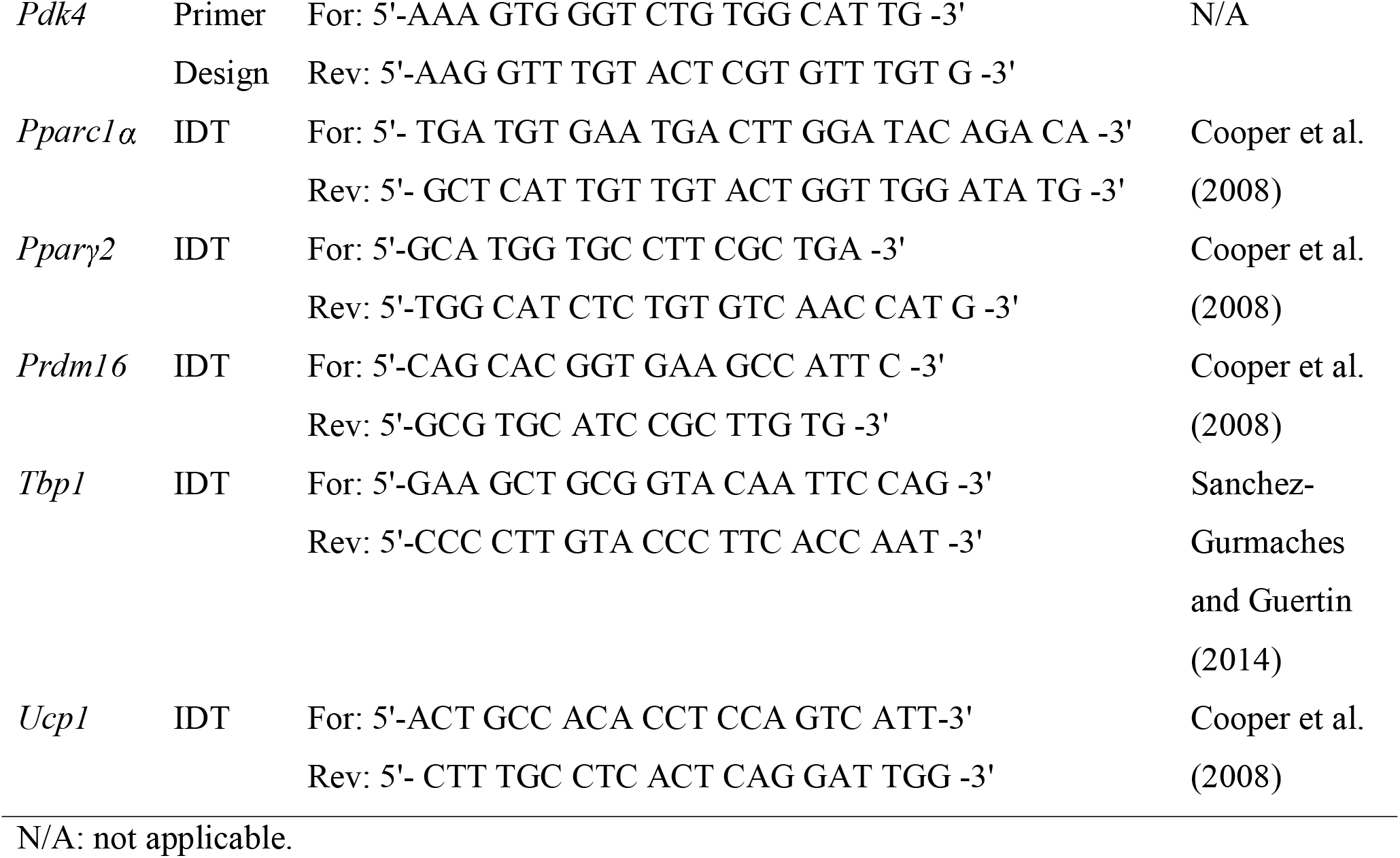
qPCR primer information.

## Acknowledgments

The authors thank Phuong Le and Casey Doucette for the assistance during the cold housing experiment, and Audrey Bergeron for critical review of the manuscript. This work was supported by the National Institute of Arthritis And Musculoskeletal And Skin Diseases (NIAMS) and the National Institute of General Medical Sciences (NIGMS) of the National Institutes of Health (NIH) under award numbers K01AR067858 and P20GM121301 to KJM. This work utilized services of the Maine Medical Center Research Institute (MMCRI) Molecular Phenotyping Core, which is supported by NIH/NIGMS P30GM106391, the Physiology Core, which is supported by NIH/NIGMS P30GM106391 and P20GM121301, the Histopathology and Histomorphometry Core, which is supported by NIH/NIGMS P30GM106391, P20GM121301, and P30103392, and the Mouse Transgenic and In Vivo Imaging Core which is supported by NIH/NIGMS P30GM103392. All cores also received support from the Norther New England Clinical and Translational Research Network NIH/NIGMS U54GM115516. The content is solely the responsibility of the authors and does not necessarily represent the official views of the National Institutes of Health.

## Authors Contributions

ALC was responsible for data acquisition, data analysis, interpretation, drafting and critical review of the manuscript. AT was responsible for data acquisition, data analysis, interpretation, drafting and critical review of the manuscript. DJB was responsible for data acquisition, data analysis, interpretation, drafting and critical review of the manuscript. SC was responsible for data acquisition, data analysis, interpretation, drafting and critical review of the manuscript. RN was responsible for data acquisition, data analysis and critical review of the manuscript. MRR was responsible for data interpretation and critical review of the manuscript. MLB was responsible for data interpretation and critical review of the manuscript. KJM was responsible for experimental design, data acquisition, data analysis, interpretation, drafting, and critical review of the manuscript. All authors had final approval of the manuscript.

