## Supplementary figures and images for "*Trpm8* modulates thermogenesis in a sex-dependent manner without influencing cold-induced bone loss"

### Supplemental Figure 1

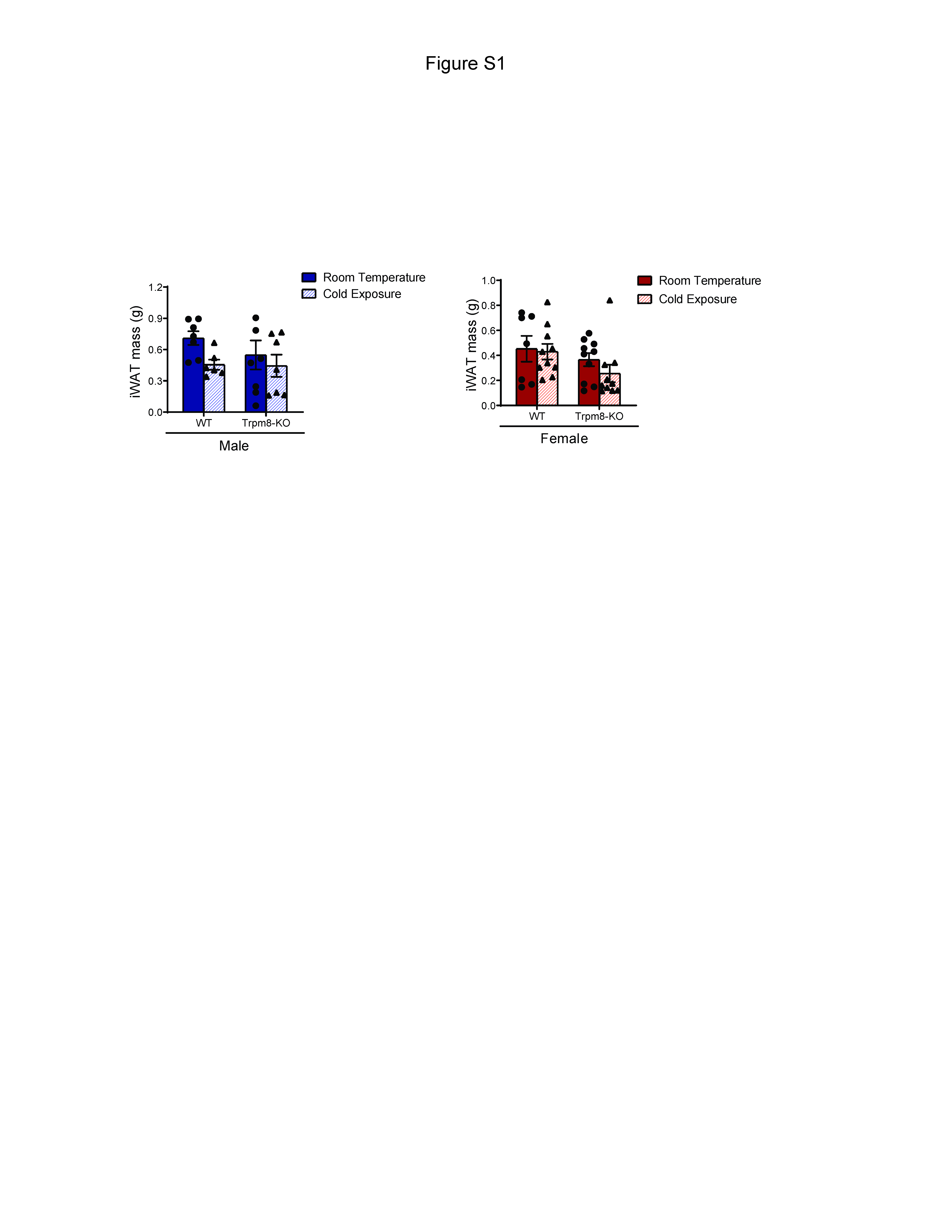
